# Faster action reprogramming, but not stopping, with proactive cues: Combining EMG and computational modelling in response-selective stop signal tasks

**DOI:** 10.1101/2023.07.11.548624

**Authors:** Sauro E. Salomoni, Quentin F. Gronau, Andrew Heathcote, Dora Matzke, Mark R. Hinder

**Author notes:** **Corresponding author:** Sauro Salomoni, Sensorimotor Neuroscience and Ageing Research Laboratory, School of Psychological Sciences, University of Tasmania,. Both authors contributed equally (Sauro E. Salomoni, Quentin F. Gronau).

## Abstract

The ability to stop simple ongoing actions has been extensively studied using the stop signal task, but less is known about inhibition in more complex scenarios. Here we used a task requiring bimanual responses to go stimuli, but selective inhibition of only one of those responses following a stop signal. We assessed how proactive cues affect the nature of both the responding and stopping processes, and the well-documented “stopping delay” in the continuing action following successful stopping. In this task, estimates of the speed of inhibition based on a simple-stopping model are inappropriate, and have produced inconsistent findings about the effects of proactive control on motor inhibition. We instead used a multi-modal approach, based on improved methods of detecting and interpreting partial electromyographical (EMG) responses and the recently proposed SIS (*simultaneously inhibit and start*) model of selective stopping behaviour. Our results provide clear and converging evidence that proactive cues reduce the stopping delay effect by slowing bimanual responses and speeding unimanual responses, with a negligible effect on the speed of the stopping process.

## Introduction

Response inhibition is an essential component of human motor control typically studied using the stop signal task (SST). Such tasks require fast responses to a “go” stimulus; a stop signal is presented on a minority of trials, requiring cancellation of the planned or initiated action. Response time (RT) to the go stimulus is directly observable, but the time required to inhibit the action (stop signal reaction time, SSRT) can only be estimated indirectly, using either non-parametric ^1^ or parametric ^2^ cognitive-modelling approaches. In contrast to the standard SST using a simple choice response, *response*-*selective* SSTs require a multicomponent response (e.g., bimanual button press) to the go stimulus, with the stop signal necessitating withholding only one component (e.g., left button press) while the other (e.g., right button press) continues to be executed as quickly as possible ^3,4^.

Macdonald et al. ^5^ proposed that response-selective inhibition is achieved through two distinct processes triggered by the stop signal: *global inhibition*, which serves to cancel the initiated multicomponent action, and initiation of the (selected) single-component response. This causes delays in RTs during successful stop trials compared to RTs in the standard multicomponent go trials, a phenomenon known as the *stopping delay* ^4,6,7^. This explanation is supported by transcranial magnetic stimulation (TMS), which indicates reduced corticomotor excitability (measured via motor evoked potential, MEP, amplitude) around 100-200 ms after the stop signal in muscles required to stop the action ^8,9^, in muscles required to continue the ongoing action component ^10^, and even in muscles not involved in the task ^11,12^. The new response is able to overcome this general motor suppression by means of a selective facilitatory input, demonstrated by larger MEPs ∼50 ms prior to the unimanual response ^10^, in addition to increased intra-cortical facilitation and reduced intra-cortical inhibition in the responding effector ^13,14^. Taken together, these findings suggest that response-selective inhibition occurs via global inhibition of all ongoing actions and the initiation of a new response, rather than a continuation of one component of the original response. Importantly, rather than a sequence of distinct motor processes, both global stopping and the new selective response are triggered simultaneously upon presentation of the stop signal, with the new response being faster and stronger than the original multi-component go response ^5,15^. As such, stopping delay arises not because of delays in the original go response, but because the new selective response has a later starting point, the stop signal.

Cognitive-control tasks, such as the SST, depend on a balance of *reactive* control and *proactive* processes based on prior knowledge or cues ^16^. *Reactive* processes dominate when the stop signal is unexpected ^17^, whereas *proactive* processes become more dominant when the potential need for action cancellation is known in advance ^18,19^. Foreknowledge may be obtained implicitly based on participants’ perceptions of the probability that a stop signal will appear in any particular trial ^20^, or explicitly by provision of contextual cues about the upcoming trial, either as the probability of a stop signal occurring ^21^ and/or which effector should be stopped ^4^. In simple stopping, most participants use foreknowledge to proactively slow their go responses in an attempt to increase the chance of successful inhibition ^21–23^. Studies investigating response-selective SSTs generally report that proactive cueing reduces the stopping delay ^4,9,24,25^. However, given proactive slowing of go-RTs, which are used as a reference to quantify the stopping delay, it is difficult to ascertain the specific effects of cueing on selective-stop trials. Moreover, it is unclear how proactive cueing affects the speed of the inhibitory process, usually assessed by SSRT, with different studies reporting that proactive SSRTs may be shorter ^26,27^, longer ^4,28^, or no different ^29,30^ than SSRTs in non-cued reactive conditions. These conflicting findings may be explained by SSRT calculations being based on a race between a go and a stop process, a model appropriate for simple stopping but clearly inappropriate when a third process (i.e., the non-cancelled action component) joins the race ^15^.

An alternative approach uses electromyography (EMG) to obtain single-trial estimates of stopping latency (for a review, see ^31^). Partial or covert EMG bursts are motor actions initiated in response to a “go” stimulus but cancelled before generating an overt button press, which can provide important information about the process of action cancellation at the single trial level. The latency of the peak of a partial EMG burst represents the moment when muscle activity associated with the initial go action starts to decrease in response to a stop signal ^31^. As such, EMG measures of stopping latency may be more closely related to neural inhibitory processes than behavioural responses (namely SSRT), which are affected by the electromechanical delay.

We recently developed a multi-modal approach that offers new insights into response-selective inhibition using both EMG measures and cognitive modelling ^15^. Our fine-grained EMG algorithm extends previous methods by detecting both partial and RT-generating EMG bursts *within the same hand* while imposing time constraints to ensure that only partial bursts related to action cancellation are captured and interpreted. Moreover, our Bayesian hierarchical *simultaneously inhibit and start* (SIS) model extends the standard race model to the response-selective SST, and so provides more plausible SSRT estimates. Here we apply this multi-modal approach to characterise the effects of proactive cues on the go and stop processes in response-selective SST, which requires bimanual response to go stimuli, but selective inhibition of one component (unimanual response) following a stop signal. Based on previous studies, we expect that proactive cues will slow down bimanual responses ^3,4^, increase the speed of unimanual responses ^29^, and reduce the stopping delay ^4,25^ compared to reactive trials. However, the effects of cues on the speed of the global stopping process are not yet clear, given conflicting reports among previous studies.

## Results

### Experimental procedures

Thirty-seven healthy adults performed *reactive* and *proactive* versions of a bimanual response-selective SST (see Figure 1). During the reactive condition, a fixation cross was presented before the bimanual go (bi-go) stimulus which, on 1/3 of trials, was followed by a stop stimulus requiring unimanual (selective) stopping, i.e., cancellation of the left or right response while continuing with the contralateral response. During the proactive condition, the fixation cross was replaced by a cue that informed participants which response would have to be cancelled (maybe stop left, maybe stop right) if indeed a stop signal was presented (1/3 trials). The stop signal delay (SSD) was staircased independently for proactive and reactive conditions to yield stop success rates of approximately 50%.

**Figure 1.**
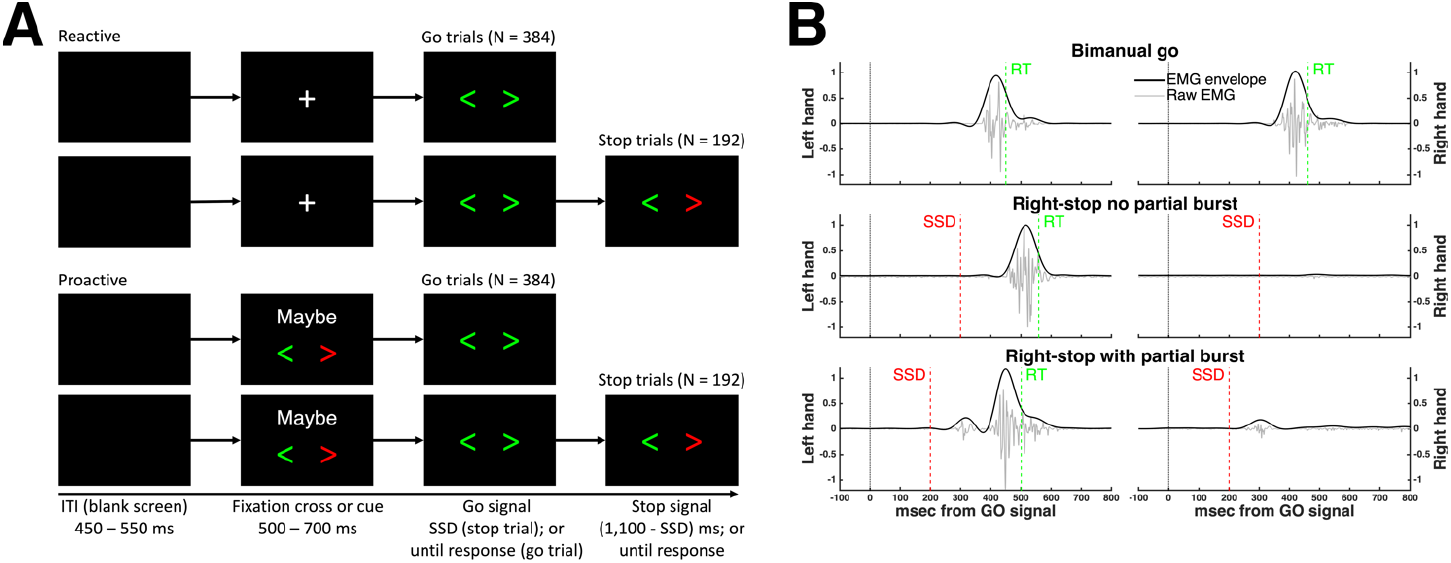
**A**. Stimuli for reactive and proactive versions of the response-selective stop signal task. A blank screen was displayed during a variable inter-trial interval (ITI), followed by a either a fixation cross (reactive) or cue (proactive) and then the bimanual go signal. In 30% of trials, the bimanual go signal was followed (after a delay of SSD ms) by a response-selective stop signal; in the proactive condition the hand required to stop was always congruent with the cue given. **B**. Representative examples of normalised EMG profiles during successful trials: Bi-go (top), selective (unimanual) stop with no partial burst (middle), and with a partial burst (bottom). Each plot shows the raw EMG, used to detect EMG onset and offset, and the corresponding EMG envelope, used to extract all other amplitude and time measures. [Figure 1]

### EMG recordings

Electromyographic signals were recorded bilaterally from the first dorsal interossei (FDI) using a belly-tendon montage with a ground electrode on the ulnar process. Within each trial, EMG burst onset and offset times were determined from the raw signals using a single-threshold algorithm ^32^, whereas EMG envelopes (full-wave rectified and low-pass filtered at 10Hz) were used to extract all other EMG measures. To allow comparisons across participants and conditions, EMG profiles from each participant were normalised to the average of the peak amplitudes across all successful bi-go trials in the reactive condition of that participant. Then, for each trial and each hand, up to two EMG bursts were detected: (i) The RT-generating burst, which resulted in the overt button press; and (ii) a partial burst, corresponding to motor responses that were initiated but cancelled prior to generating sufficient force to register a button press. Because we only expect responses to be cancelled after presentation of a stop signal, partial EMG bursts were only assessed in successful stop trials. Importantly, we imposed timing and amplitude constraints for the detection of partial bursts that prevent inclusion of spurious muscle activity unrelated to the stop signal task, such as mirror activity or other spurious voluntary or involuntary activation unrelated to the task. Briefly, an EMG burst is only considered to represent a partial response if it occurs between SSD and the onset of the RT-generating burst, in addition to having amplitude > 10% of the average EMG amplitude observed in reactive bi-go trials (for details, see Methods section).

### Effects of proactive cues on behavioural and EMG measures

Proactive slowing was evident as longer bi-go RTs in the proactive vs. reactive condition (Figure 2, interaction Condition * Trial Type * Success, *p* < 0.001) both during successful bi-go trials (proactive vs. reactive: 480 ± 3 ms vs. 451 ± 3 ms) and failed selective stop trials (424 ± 5 vs. 394 ± 5 ms). As predicted by the standard race model and SIS, failed stop-RTs were shorter than successful go-RTs for both conditions (both *p* < 0.01). In contrast, successful stop-RTs were longer than go-RT for both conditions (stop-RT > go-RT proactive: 555 ± 5 ms > 480 ms, reactive: 554 ± 5 ms > 351 ms, *p* = 0.01), consistent with the stopping delay reported in previous studies ^3,4,25^. We also found that the magnitude of the stopping delay (interference effect) was reduced in the proactive vs. reactive condition (68 ± 10 ms vs. 97 ± 7 ms; *p* < 0.001).

**Figure 2.**
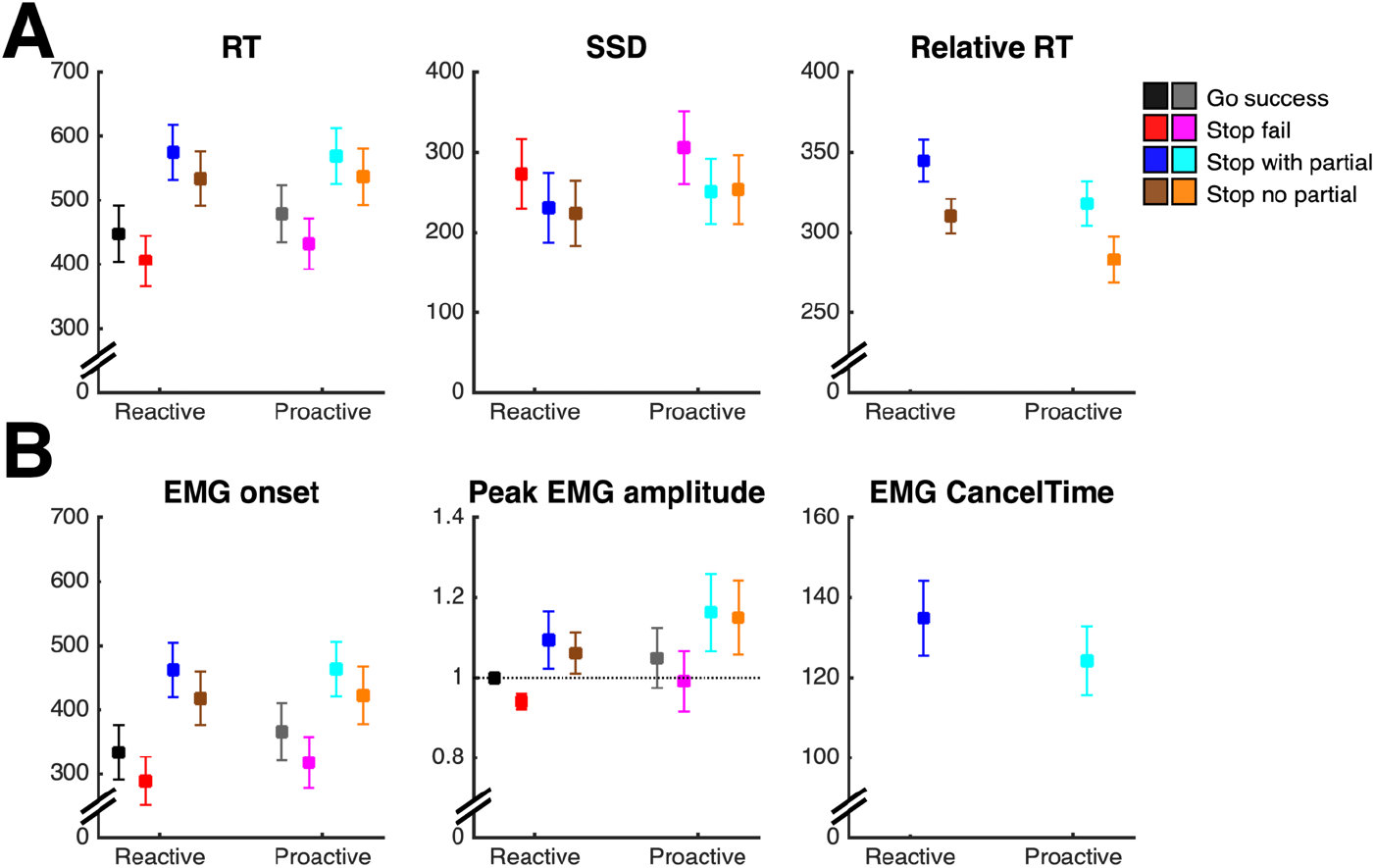
**A**. Behavioural measures (average ± 95% CI) in the proactive and reactive conditions, as a function of the type of response: Successful bi-go, failed stop, successful stop with and without a partial burst. Stopping delay is evident in both conditions as longer stop-RT than go-RT. During successful stop trials, the selective (unimanual) response is by SSD (relative RT), with shorter stop-RTs in proactive than reactive condition. **B**. EMG measures (average ± 95% CI). Note that the peak EMG amplitude of each subject was normalised to the average amplitude during successful reactive bi-go trials (marked by dotted line), hence the small error bars in reactive bi-go and failed stop trials, which tend to have similar amplitudes.

As a direct consequence of proactive slowing, SSD was longer in the proactive vs. reactive condition (267 ± 5 ms vs. 238 ± 5 ms, interaction Condition * Success, *p* = 0.004), although SSD staircasing ensured similar stopping success rates across both proactive (49.5%) and reactive (47.8%) conditions. To assess selective (unimanual) stop-RTs, we need to consider when this process is initiated. Consistent with model frameworks for response-selective stopping ^5,15^, we assume that motor commands for the bi-go and unimanual (non-cancelled) selective responses are initiated at different time points: The bimanual response is initiated upon presentation of the go signal, whereas the selective response is initiated after the (selective) stop signal (i.e. SSD ms after the go signal). When measuring successful stop-RT relative to SSD, we found that this “relative RT” was substantially faster than the bi-go RT for both proactive (289 ± 3 ms) and reactive conditions (316 ± 3 ms, interaction Condition * Success, *p* < 0.001). When compared between conditions, relative RTs were shorter in the proactive compared to the reactive condition (*p* < 0.001), indicating faster selective (unimanual) responses because of proactive cueing.

Analyses of EMG timings are consistent with our behavioural results: During successful bi-go trials, the onset of the RT-generating burst occurred later during proactive than reactive condition (Figure 2, proactive vs. reactive: 359 ± 3 ms vs. 328 ± 3 ms; interaction Condition * Trial Type * Success, *p* < 0.001), supporting the notion of proactive slowing. In both conditions, the behavioural stopping delay was also confirmed by later EMG onset times in successful stop trials than in bi-go trials (EMG onset stop > bi-go proactive: 442 ± 3 ms > 359 ms; reactive: 438 ± 3 ms > 328 ms; both *p* < 0.01). Given our assumption that the new unimanual response is triggered by the stop signal, we measured EMG onset relative to SSD and we found that the relative EMG onset occurred earlier in proactive than in reactive trials (182 ± 2.7 ms vs. 207 ± 2.5 ms; interaction Condition * Success, *p* < 0.001).

Averaged over all trial types and success outcomes, no significant main effect of Condition was observed in the (normalised) amplitude of the RT-generating burst; however, there were significant interactions between Condition * Trial Type and between Condition * Success. Across both conditions, the peak EMG amplitude was lower during failed stops (0.96 ± 0.01) than during successful bi-go trials (1.02 ± 0.01, interaction Trial Type * Success, *p* < 0.001), suggesting an attempt (albeit unsuccessful) to inhibit the ongoing action. In contrast, during successful stop trials, the amplitude of the re-initiated response (1.12 ± 0.01) was higher than both failed stop trials and successful bi-go trials (both *p* < 0.01). These results suggest that, for both proactive and reactive conditions, the new unimanual response, triggered upon presentation of the stop signal, is not only faster, but also stronger than the initial bimanual response consistent with Gronau et al. ^15^. Proactive cues caused a reduction in the proportion of successful stop trials in which a partial burst was detected (proactive vs. reactive: 15.0 ± 1.1% vs. 19.9 ± 1.4%, main effect of Condition, *p* < 0.001), suggesting greater efficiency of stopping (i.e., motor activity was more completely cancelled) when proactive cues were provided. Finally, we found a very small, albeit significant, reduction in the latency of the peak of the partial burst relative to SSD (EMG CancelTime proactive vs. reactive: 122 ± 4 ms vs. 130 ± 4 ms; interaction Condition * Success, *p* = 0.008), but the small absolute difference (8ms) and effect size (Cohen’s d = 0.17) suggests the effect of this difference is negligible.

We used statistical parametric mapping (SPM1d ^33^) to compare EMG profiles between the proactive and reactive conditions (see Figure 3). SPM indicated that, in trials without a partial burst, the RT-generating burst was initiated earlier in the proactive than reactive condition, supporting our behavioural and model-based findings indicating faster unimanual responses in successful selective stop trials (relative to SSD) when proactive cues were provided. When EMG profiles were aligned to the stop signal, SPM revealed no significant differences in the shape or timing of the partial (cancelled) EMG burst between proactive and reactive conditions, suggesting similar stopping latencies (CancelTime) of the initial bimanual response across both conditions.

**Figure 3.**
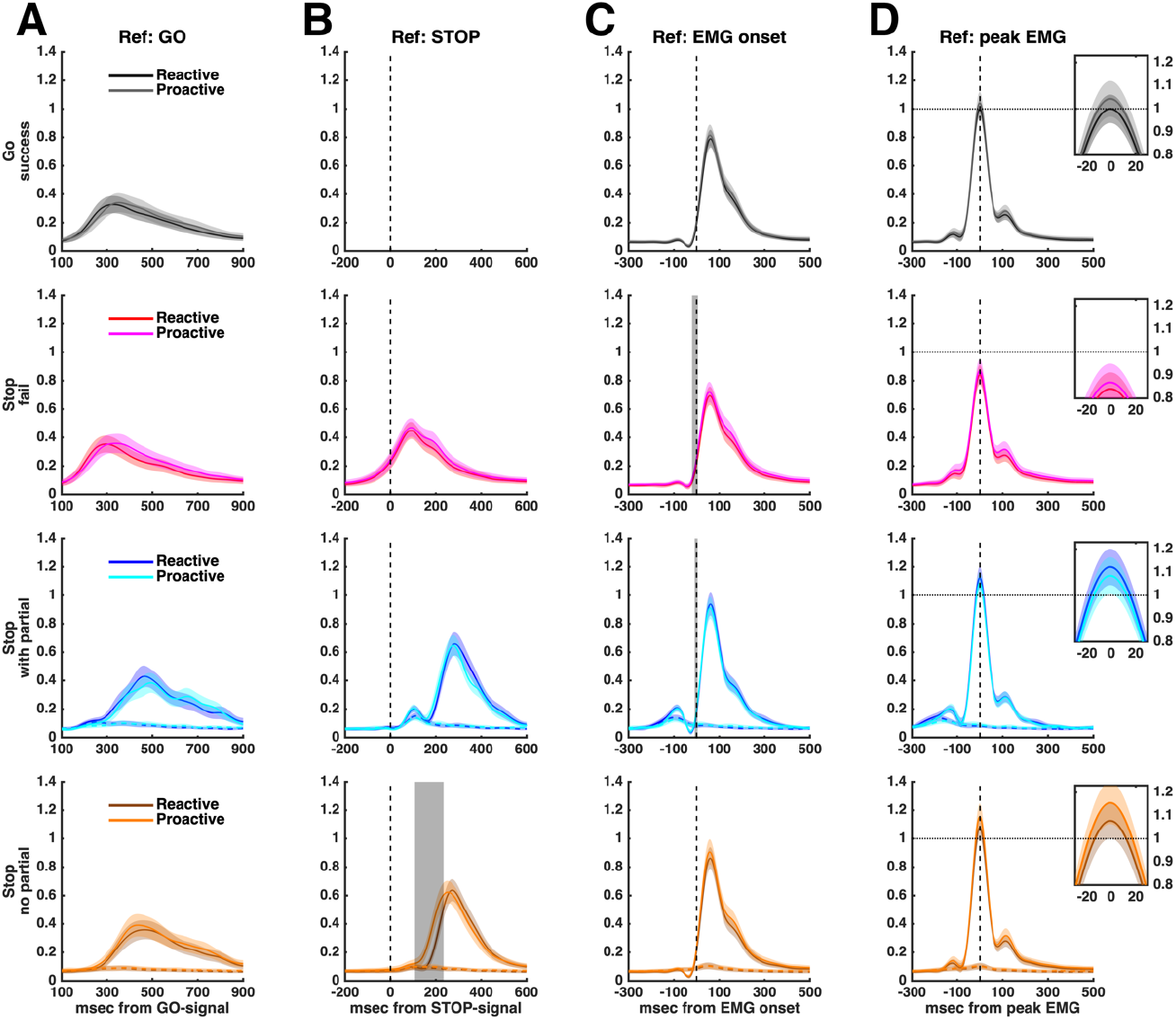
EMG profiles, showing the average ± 95% CI across subjects aligned to the presentation of the go stimulus, stop stimulus, EMG onset and peak EMG (columns **A**-**D**, respectively). Successful stop trials were divided in those with or without a partial burst (3^rd^ and 4^th^ rows). EMG amplitude was normalised to the average amplitude of successful bi-go responses in the reactive condition (horizontal dotted line in **D** Ref: peak EMG). The averaged EMG profiles have smaller amplitudes when shown in reference to the go and stop signals (**A** and **B**) or EMG onset I, as the individual peaks are not coincident. Shaded grey areas indicate periods where EMG profiles differ significantly between proactive and reactive conditions, based on SPM analyses (see Methods). In particular, the bottom row (Stop no partial) in **B** indicates that in successful selective stop trials without a partial burst, the unimanual response occurs earlier (relative to the stop signal) in the proactive than in the reactive condition. In **D**, the small ‘zoom’ inlet plots of peak EMG show that, for both conditions, the amplitudes of failed stop responses (2^nd^ row) are lower, whereas successful stop responses (3^rd^ and 4^th^ rows) are greater, than the amplitude of go responses (1^st^ row).

Similar to our previous work ^15^, we used the full EMG time series from all successful stop trials of all participants to investigate the presence and timing of partial and RT-generating bursts (Figure 4). Critically, Figure 4 indicates a continuous distribution of timing between the two bursts, suggesting a temporal merging of the cancelled bimanual response and the subsequent unimanual response. Specifically, the delay from the offset of the partial burst to the onset of the RT-generating burst followed a continuum from 273 ms to 20 ms, with a distribution skewed towards the shorter time, where the lower bound of 20 ms corresponds to the limit of our EMG detection algorithm for the detection of two distinct bursts. Beyond the limit of our algorithm, when we could only detect a single burst, the longer “tail” on the left side of the single burst occurs at a similar latency as the latest partial bursts, suggesting a temporal merging of the two processes, i.e., cancellation of the initial bimanual action and initiation of a unimanual response. This observed merging of partial and RT-generating bursts is wholly consistent with the framework of our computational model. Specifically, the (global) stop process does not need to be completed (i.e., win the race to threshold) before initiation of the response-selective (unimanual) response: If that were the case, we should observe a finite delay between the partial burst (bimanual stop) and the RT-generating burst (unimanual go) rather than continuous, overlapping distributions (Figure 4 **C** and **D**). Instead, our results strongly suggest that both the bimanual stop and unimanual go processes are simultaneously triggered at SSD, with the presence and timing of partial bursts determined by the outcome of the “race” between the bimanual go and stop processes.

**Figure 4.**
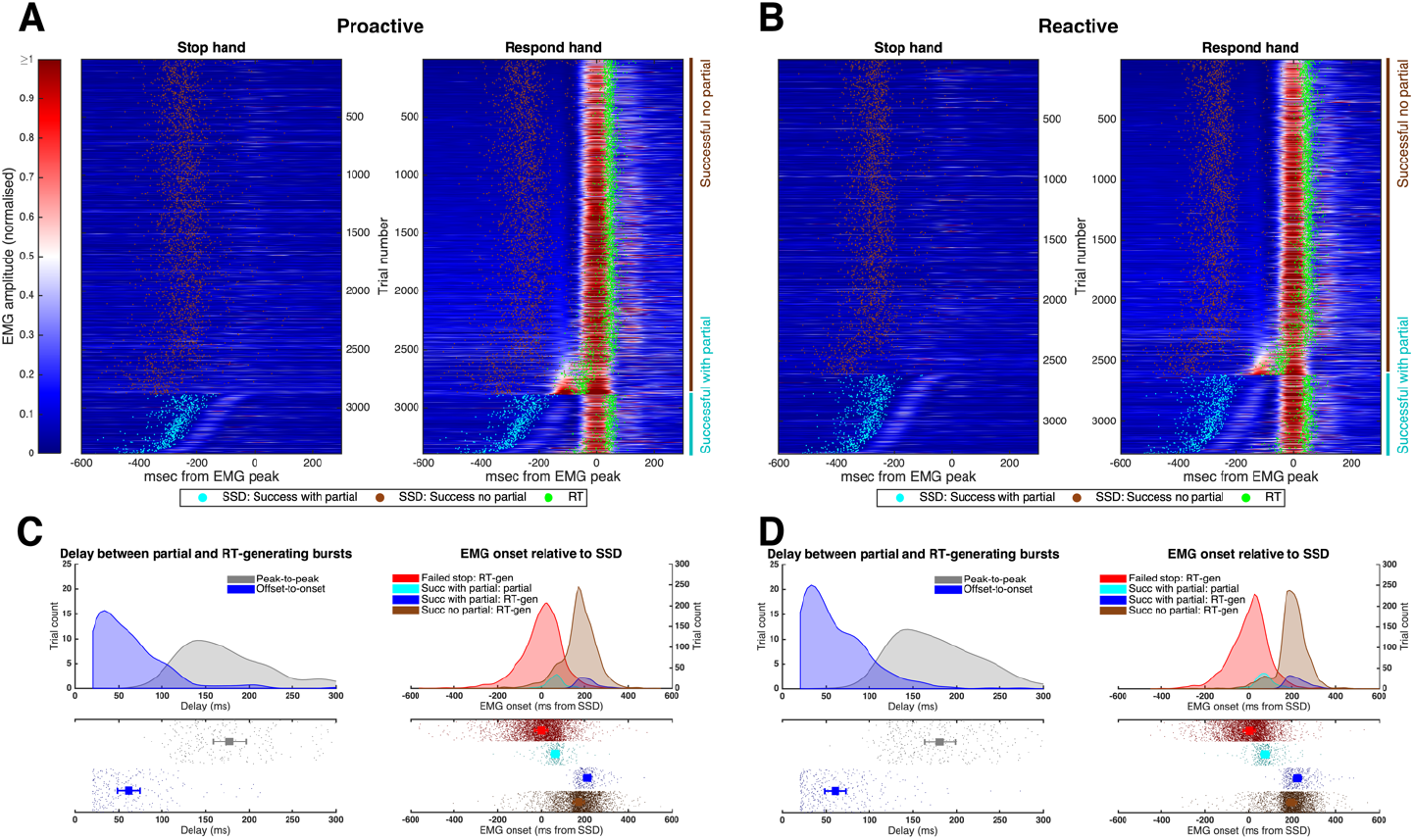
Single-trial EMG profiles from all successful stop trials (pooled across all subjects) during (**A**) proactive and (**B**) reactive conditions, respectively. The EMG signals from each trial were colour-coded by the normalised amplitude and referenced to the peak of the RT-generating burst. Trials with a partial burst (in either hand) are displayed at the bottom of each panel, sorted by the delay between the partial and RT-generating bursts. The remaining trials, without a partial burst, were sorted by the EMG amplitude prior to the RT-generating burst. Histograms in **C** (proactive) and **D** (reactive) show the corresponding delay between partial and RT-generating bursts, represented as peak-to-peak or offset-to-onset times, in addition to the time of EMG onset of both the partial and RT-generating bursts relative to SSD. These results suggest a temporal “merging” between the partial and RT-generating bursts, consistent with our SIS model framework.

### Modelling: Simultaneously inhibit and start (SIS)

The SIS model extends the architecture of the classic race-model ^2^ to account for response-selective stopping ^15^. It assumes that a selective stop signal simultaneously inhibits the initial bimanual response and triggers a new selective (unimanual) response. Here, we applied the SIS model in a Bayesian hierarchical manner to fit and compare data from proactive and reactive conditions (Figure 5). We report four variations of the model, where each of the go and stop processes was either allowed to vary or was fixed across proactive and reactive conditions (for details, see Methods section). Assessment using Ando’s BPIC criterion ^34^ favoured Model 1 (BPIC = 59860.74), in which parameters for the bi-go and unimanual left/right runners were allowed to vary with cueing condition, but parameters for the dual-stop runner were invariant across conditions.

**Figure 5.**
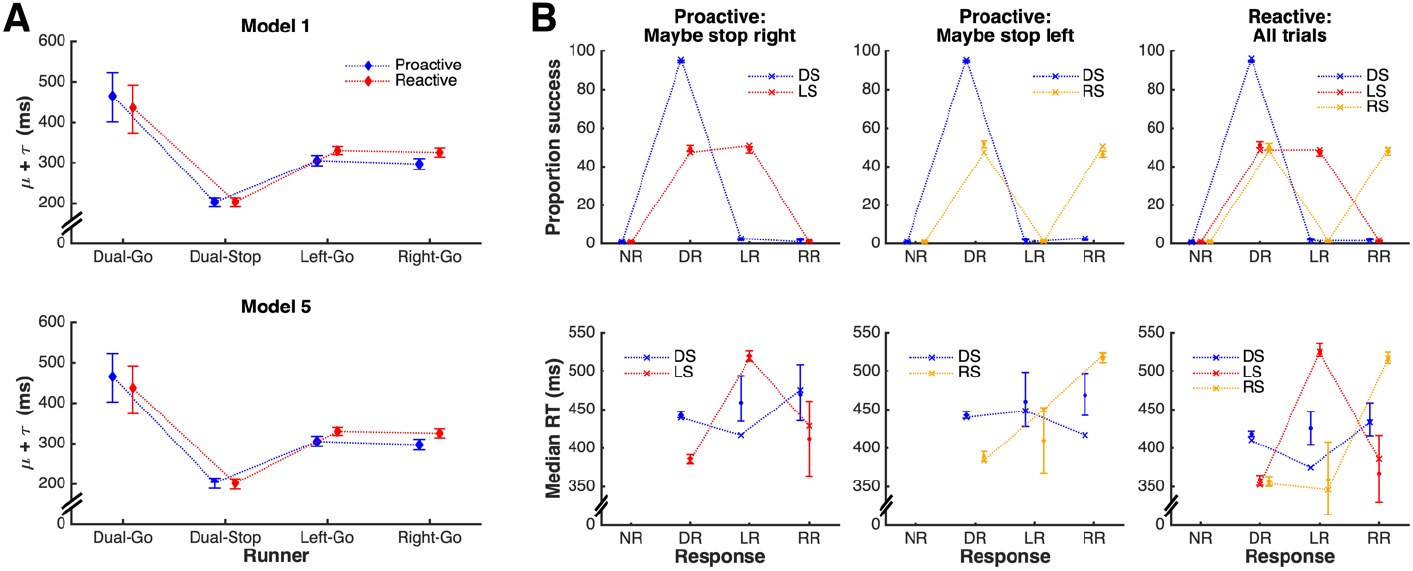
**A**. Estimated running times (i.e., *μ + τ* estimates) for the dual-go, dual-stop, and left/right selective runners in the SIS model, expressed as the posterior median for this quantity (in ms), accompanied by the 95% credible interval. **Model 1** corresponds to the best estimates based on BPIC, in which the dual-stop runner was constrained to have the same running time in both the proactive and reactive conditions. **Model 5** corresponds to the second-best model in terms of BPIC, which relaxed this constraint. It is apparent that the estimates are very similar for both models. Importantly, even when the dual-stop runner is allowed to vary across conditions (**Model 5**), individual-level estimates (not displayed) do not suggest a meaningful difference (median difference: 1 ms, 95% credible interval [-2 ms, 5 ms], see Table 1). **B**. Model fits for the various combinations of stimulus (DS: Dual stimulus, LS: Left stimulus, i.e. right-stop, RS: Right stimulus, i.e. left-stop) and response (NR: No response, DR: Dual response, LR: Left response, RR: Right response) for the model with the best fit (**Model 1**). Data is shown as crosses connected by dotted lines, with model fits displayed as median across posterior predictions, accompanied by 95% credible intervals. The model provides a good representation of the data, given that estimates show a close fit to data with tight credible intervals for all common cells. Signs of misfit on median RT are only apparent for very rare responses, e.g., left response to a dual stimulus; left response on a right stimulus (left-stop) trial.

Inspection of the parameter estimates reveals that proactive cueing resulted in *slowing* of the dual-go runner, consistent with the behavioural data that exhibit the typical proactive slowing. Examination of the difference between the cueing conditions revealed this slowing was non-negligible (Table 1, median difference and 95% credible interval: 28 [26, 30] ms). In both cueing conditions, the speed of both unimanual runners was substantially faster than the speed of the dual-go runner in the corresponding condition; differences were of the order of 120 ms in the reactive condition and ∼175-180 ms in the proactive condition. Finally, the speeds of both the left and right unimanual runners were faster in the proactive compared to the reactive condition (median difference and 95% credible interval left -25 [-28, -22] ms, right -28 [-31, -25] ms), consistent with our behavioural measure of relative RT.

**Table 1.** SIS model individual-level estimates: Contrasts of interest (in milliseconds). Estimates obtained using the model where the stop runner was allowed to vary between conditions (using parameters from Model 5, second-best fit).

| Contrast | Median | 95% Credible interval |
| --- | --- | --- |
| Proactive - Reactive |  |  |
| Proactive Dual-Go - Reactive Dual-Go | 28 | [26, 30] |
| Proactive Left-Go - Reactive Left-Go | -25 | [-28, -22] |
| Proactive Right-Go - Reactive Right-Go | -28 | [-31, -25] |
| Proactive Stop - Reactive Stop | 1 | [-2, 5] |
| Dual-Go runner - Unimanual-Go runner |  |  |
| Proactive Dual-Go - Proactive Left-Go | 173 | [171, 176] |
| Proactive Dual-Go - Proactive Right-Go | 181 | [179, 184] |
| Reactive Dual-Go - Reactive Left-Go | 120 | [117, 123] |
| Reactive Dual-Go - Reactive Right-Go | 125 | [122, 128] |

Interestingly, the second-best model (?BPIC = 16.4 compared to Model 1) was the one in which the dual-stop runner’s parameters *were* permitted to vary by cueing condition. Critically however, despite this flexibility, individual-level estimates do not suggest a meaningful difference in dual-stop parameters between the proactive and reactive conditions (median difference and 95% credible interval: 1 [-2, 5] ms). These findings strongly suggest that cueing has a negligible effect on the stop process, but rather significantly affects the speed of the dual-go and unimanual go processes. The two other model variants, which constrained the dual-go or unimanual runners across conditions, are extremely unlikely to represent the data (?BPIC > 900 and > 3600 compared to Model 1, respectively).

We used the model with the best fit to further investigate the small difference between conditions found in EMG CancelTime (Figure 2; for details, see supplementary material). In trials with a partial EMG response, the finish times of the stop runner correspond approximately to EMG CancelTime, and the finish time of the unimanual go runner corresponds to stop-RT. Given that estimation of EMG CancelTime requires the presence of an observable partial EMG burst, it excludes trials where stopping is “too fast” (no partial burst) or “too slow” (either failed stops, or where the partial burst is so close to the RT-generating burst that it cannot be detected) – for a similar discussion, see Figure 4 in Raud et al. ^31^. We incorporated these limits in the model simulations by imposing lower and upper thresholds on how much earlier the finish time of the stop runner must be relative to that of the dual-go runner. The upper threshold (stop is “too slow”) was conservatively estimated as 90 ms, comprising the sum of (i) time from peak to offset of the partial burst (∼50 ms); (ii) minimum period from offset to onset (20 ms); and (iii) time from onset of the RT-generating burst to the unimanual button press, i.e., stop-RT (∼20 ms, excluding the electro-mechanical delay). We estimated the lower threshold (“too fast”) as 115 ms, as this results in a proportion of trials with partial burst that is consistent with our data (∼18% of successful stops). Discarding (censoring) any trials beyond these two thresholds resulted in down-bias of the estimated finish times of the stop runner (i.e., faster speed) in both conditions. This down-bias was ∼7 ms stronger in proactive than reactive (credible interval of difference: 5 – 9 ms), causing the stop runner to be faster in proactive, which is wholly consistent with the 8 ms difference found in EMG CancelTime. In further investigations, we varied the upper and lower thresholds across a range of reasonable values and generally found a down bias occurred often, although it was sometimes of a smaller magnitude. Although approximate, this simulation clearly suggests that the small difference found in EMG CancelTime results from an artifact of the censoring inherent to this measure of stopping latency.

In summary, we provide converging evidence from behaviour, EMG, and cognitive modelling indicating that during a response-selective stop signal task, proactive cues result in slowing of the initial bi-go responses and increased speed of the new selective (unimanual) response, both of which contribute to a reduction in, but not elimination of, the extent of the stopping delay. Critically, our results strongly suggest negligible effects of proactive cues on the speed of the stopping process.

## Discussion

Using a multimodal approach combining electromyographical recordings and Bayesian computational modelling, the current study provides novel insights into how proactive cues can modify performance of a response-selective stop signal task. Specifically, using our recently developed SIS model and fine-grained analysis of both covert and overt EMG responses ^15^, we were able to tease apart the effect of proactive cueing on the inhibitory process and on the ongoing action components. Overall, we observed that proactive cues resulted in significant *slowing* of the bimanual response, whereas the unimanual response in successful stop trials was significantly *faster* in this proactive condition. Together, these cue-related changes in motor action resulted in a substantial reduction in the well documented stopping delay ^25^. In contrast, changes in the speed of the rapid inhibitory process because of proactive cueing were either absent or negligible. Specifically, both modelling results and statistical parametric mapping (SPM) suggest no change in the inhibitory process. Nevertheless, the smaller *proportion* of successful stop trials in which a partial EMG response was observed suggests that stopping *efficiency*, rather than stopping *speed*, did improve as a result of proactive cueing.

### Distinct effects of proactive cueing on bimanual and unimanual responses

Our results replicate previous studies reporting that proactive cues slow go responses compared with reactive (non-cued) trials, which has been demonstrated in both simple and response-selective versions of the stop signal task ^4,35^. This effect is believed to represent a proactive slowing strategy attempting to maximise the likelihood of stopping success ^36^. Specifically, it has been suggested that proactive control is influenced by adjustments in attention in other (non-inhibitory) contexts ^16,37^, which have been linked to anticipatory activity in the visual cortex and other sensory areas ^38–40^. For example, fMRI studies report an association between proactive slowing of go-RTs and functional connectivity between the ventral striatum and visual areas ^41^, likely associated with processing visual cues.

In the current study, we used a response-selective variation of the stop signal task. In this task, presentation of a selective stop signal simultaneously triggers both inhibition of the initial bimanual response and initiation of a new unimanual response ^5^. Consistent with our previous work ^15^, we found that the newly programmed selective response was faster and stronger than the initial bi-go response, for both cued and non-cued conditions. Compared between conditions, the responses to selective stop trials were faster in the proactive condition, when participants had knowledge about which response may have to stop (and thus which response would have to continue) in the upcoming trial. Studies using EEG have shown considerable functional overlap between preparation for withholding a planned response and switching it with a different response ^42^. The result is a highly flexible proactive control system that can be tailored to the expected goals on a trial-by-trial basis to selectively target a particular response ^19,21^, such as changing from bimanual to unimanual response. For example, sustained intra-cortical inhibition has been observed during simple action preparation, whereas task switching was associated with a release of this inhibition, suggesting that visual cues may influence primary motor cortex output in anticipation of movement initiation ^43,44^. In our study, although stop signals comprised only 30% of trials and were therefore unexpected, modulation of attention involved with the visual cues may have proactively activated the motor network, biasing action selection and increasing the speed of the alternative response ^45^.

### Proactive cueing results in more efficient, but not faster, stopping performance

We observed multiple lines of evidence to suggest that inhibitory control in this response-selective stopping task is more efficient when proactive cues indicating which effector may have to countermand its response are provided. First, the proportion of successful stop trials in which a partial EMG burst was observed was smaller in proactive compared to the reactive condition, indicating that the bimanual response was more often inhibited before any detectable muscle activity had been initiated. Second, the stopping delay (interference effect) observed during successful selective stop trials was reduced, but not abolished, by proactive cues (for a recent review, see ^25^).

Recent evidence suggests that the stopping delay emerges because of non-selective response inhibition during selective stop trials, whereby both the effector that is required to stop as well as the effector required to continue its action are inhibited before a new response can be initiated ^46^. This non-selective inhibition is followed by a new, selective (unimanual) response, likely through a combination of disinhibition and excitation of the responding effector ^5^. Hence, the magnitude of stopping delay reflects both the delay induced by response inhibition and the speed of the new selective response.

The traditional view posits that reactive and proactive inhibition are implemented by fast global and slow selective inhibitory systems, respectively, underpinned by distinct neural pathways ^4,6^. This theoretical dichotomy is often cited as a putative mechanism to explain the smaller “cost” of stopping when proactive cues are provided, as indicated by reduced stopping delay ^25^. Although true for simple-stopping, this may not be the case in response-selective stop signal tasks. Functional neuroimaging studies suggest that selective stopping may involve both the hyperdirect *and* indirect pathways ^9,12,47^, with the balance between these two shifted by the extent of proactive control.

Corroborating, our findings suggest that the reduction in stopping delay in the proactive condition is not associated with differences in stopping speed, but instead results from a combination of slower bimanual responses and faster unimanual responses. These were observed in our data as differences in behavioural measures and modelling parameters of go-RT and stop-RT, in addition to SPM analyses showing that EMG responses were initiated earlier in the proactive than in the reactive condition, at least in the 80-85% of successful selective stop trials, where no partial burst was observed. These findings further support the idea that proactive cues can reduce the time required to switch from a (prepotent) bimanual to an (alternative) unimanual response, likely mediated by increased attention in anticipation of the potential need to selectively stop the cued effector ^42^.

Despite these differences in go-RT and stop-RT, converging results suggest the *speed* of inhibition was unaffected by proactive cueing. Specifically, the best model, selected on the basis of the Bayesian predictive information criterion ^34^, constrained the stop process to be invariant between proactive and reactive conditions. Moreover, even when we considered the next best model, which allowed the stop process to vary according to cueing condition, contrasts between conditions showed no significant differences in the finish times of the stop runner (see Figure 5 and Table 1 for model comparisons), providing strong evidence that the speed of the stop runner did not vary between cueing conditions. Furthermore, SPM analyses of EMG profiles also suggest no differences in the timing or amplitude of successfully cancelled (partial) responses, a proxy of stopping speed.

Although results from GLMM found that EMG CancelTime was ∼8 ms faster in proactive vs. reactive trials, the associated effect size was negligible (Cohen’s d = 0.17). Moreover, simulations from our Bayesian model suggest this difference is a direct consequence of inherent censoring associated with this EMG measure of stopping latency, and not due to differences in the neurophysiological stopping process itself. Specifically, as the measure of EMG CancelTime relies on the presence of detectable partial EMG bursts, it precludes the use of trials where the stopping process is “too fast” (no partial burst) or “too slow” (either failed stop trials, or the delay between partial and RT-generating burst is less than 20 ms, the limit of our EMG detection algorithm). We performed simulations imposing similar censoring constraints on the timing of the go and stop processes (see supplementary material for details), which caused a down-bias in the speed of the stop runner, with stronger effects in the proactive condition, resulting in estimated stop times that are ∼7 ms longer in the reactive vs. proactive trials – consistent in size and direction with the small difference observed in EMG CancelTime.

In conclusion, our multi-modal approach, based on behavioural measures, EMG and computational modelling, provides converging evidence that proactive cues can result in task adjustments that strongly affect the speed of both the initial multi-component (bimanual) response, as well as the reprogrammed selective (unimanual) response, but have negligible effects on the speed of the already fast stop process.

## Methods

### Participants

Thirty-seven healthy participants aged between 18 and 36 years (mean ± SD: 24 ± 5.25 years) volunteered for this study, which was approved by the Tasmanian Human Research Ethics Committee (protocol H0016981). Written consent was obtained from all participants prior to data collection, and the study was performed in accordance with all relevant guidelines and regulations, and with the Declaration of Helsinki ^49^. One participant was not able to complete the task accurately, and had the data removed from the analysis.

### Experimental task

Participants were seated approximately 80 cm away from a computer monitor. The forearms were pronated on a desk about shoulder width apart, with each index finger resting on a button. The experimental procedures were explained verbally before trial start, and Psychopy ^50^ was used to present visual instructions on a black screen. Each button required a finger flexion force of at least 1N in order for the response to be registered.

Each participant performed a total of 1,242 trials in a single experimental session, comprised of 1,152 experimental trials, divided between reactive and proactive conditions (576 each), in addition to 90 practice trials: 30 go-only trials before the start of the experiment, and 30 reactive or proactive trials before the start of the respective condition. At the start of each trial, a white fixation cross was presented at the centre of the computer screen for 500 – 700 ms (uniform distribution) before the go stimulus, which consisted of two green arrows pointing left and right, indicating a bimanual response requiring flexion of both the left and right index fingers. The variation in presentation period aimed to prevent temporal prediction of the upcoming go stimulus. After presentation of the go stimulus, a response time window of 1,100 ms was allowed before termination of each trial.

In 1/3 of trials, the bimanual stimulus was followed by a unimanual stimulus to either stop-left or stop-right (96 trials each). All participants performed the task in both reactive and proactive conditions. In the reactive condition, one of the green arrows turned red after a variable stop signal delay (SSD), instructing the participant to stop the hand indicated by the direction of the red arrow while continuing the button press with the contralateral (respond) hand. The SSD was initialised as 200 ms and stair-cased (independently for left- and right-stop) to ensure success rates of approximately 50%, i.e. SSD was reduced by 50 ms after a successful stop or increased by 50 ms after a failed stop ^1^.

A similar procedure was used for the proactive condition, except that the initial fixation cross was replaced by a proactive cue that was either symbolic (red left or right arrow) or textual (“Maybe stop left” or “Maybe stop right”). The proactive cue was displayed at the start of all trials, with the direction balanced between left- and right-stop, while keeping the same ratio of 1:3 between stop and bi-go trials. Both the order of conditions (reactive or proactive) and the type of proactive cue used (symbol or text) were balanced across participants.

Visual feedback was provided after each trial, displaying a different message according to task performance: (i) RT following successful trials; (ii) “failed to stop” after a bimanual response on a selective stop trial; (iii) “stopped wrong hand” in case the participant stopped the incorrect hand; (iv) “missed” if no response was detected, or (v) “please press simultaneously” if the delay between button presses was more than 50 ms in bi-go trials.

### Electromyographic (EMG) recordings

EMG signals were recorded using adhesive electrodes (Ag/AgCl) positioned in a belly-tendon montage on the first dorsal interosseous (FDI) muscle of each hand, with a ground electrode positioned on the ulnar bone on each wrist. The analogue signals were band-pass filtered at 20–1,000 Hz, amplified 1,000 times, sampled at 2,000 Hz (CED Power 1401 and CED 1902, Cambridge, UK) and saved into a PC for offline analysis. Prior to commencing the task, participants were instructed to completely relax their hand muscles. Participants were reminded to relax their hands whenever their tonic muscle activation increased during the experiment, i.e., not related to stimuli presentation.

### Data processing

Behavioural data were exported from PsychoPy and included RT and SSD from each trial. For each participant, we calculated the amount of selective stop delay as the difference between the average RT during successful selective stop trials (i.e., left-stop and right-stop) and the average RT in bi-go trials, separately for reactive and proactive conditions. Similarly, the amount of proactive slowing of each participant was quantified as the difference in average successful bimanual RTs between proactive and reactive conditions.

The EMG signals were digitally filtered using a fourth-order band-pass Butterworth filter at 20–500 Hz. The precise times of EMG onsets and offsets were detected using a single-threshold algorithm ^32^. For each trial, the segment with the lowest RMS amplitude (baseline) was identified using a moving average window of 500 ms and used to identify all EMG bursts with an amplitude of more than 3 SD above baseline. For robustness, EMG bursts separated by less than 20 ms were merged together.

From the onsets and offsets detected, we set time and amplitude constraints to identify up to two bursts of interest: First, the RT-generating burst, which caused the button press, was identified as the last burst where (i) the onset occurred after the go signal and (ii) at least 50 ms before RT. Second, we also searched for partial EMG bursts on stop trials, which represent responses that were initiated after the go signal but cancelled (in response to the stop signal) before generating a button press – also called ‘partial EMG’ or ‘partial response EMG’ ^31^. Surprisingly, however, even though partial EMG bursts are postulated to represent cancellation of the initial action, some response-selective SST studies have reported them in the stopping effector *concurrently*, or even *after*, a RT-generating burst (e.g., ^35^). Such timings are inconsistent with the notion of global inhibition followed by the initiation of a new response ^5,10^, and would require strong decoupling between effectors.

Alternatively, they likely represent a consequence of EMG detection algorithms that don’t constrain the time at which partial EMG can be detected and/or are unable to identify both partial and RT-generating bursts in the same hand. With these limitations, in addition to the true action-cancellation partial bursts, other bursts may also be detected, including mirror activity and spurious (involuntary/unrelated) EMG, therefore hindering interpretation about the timing of partial bursts solely representing action-cancellation processes. Here we imposed time and amplitude constraints to avoid EMG bursts unrelated to the stop signal task: The partial bursts were identified in each hand as the earliest burst where (i) EMG onset happens after the go signal; (ii) time of peak EMG happens after SSD (i.e., inhibition happens in response to the stop signal); (iii) time of peak EMG is earlier than the onset of the RT-generating burst; and (iv) peak EMG amplitude is greater than 10% of the average peak from successful bi-go trials.

Finally, EMG profiles were obtained by full-wave rectifying and low-pass filtering at 10 Hz. For each subject, single-trial EMG profiles from both proactive and reactive conditions were normalised using the average peak EMG amplitude across successful bi-go trials from the reactive condition. This normalisation allows direct comparison of amplitude between conditions.

### Model fitting and selection

Behavioural data were fit using our newly developed model for response-selective stopping ^15^. This model, the Simultaneously Inhibit and Start (SIS) model extends the simpler independent-race model used for standard stop signal tasks ^2^ by including a third runner, representing the response of the non-cancelled action component (e.g., the unimanual right response on a left-stop trial), which commences at the onset of the unimanual stop signal (see ^15^ for more details on the model framework). In the current task we focused solely on response-selective stopping (i.e., bi-stop trials were not present), although our model allows the inclusion of a bi-stop process, where the bimanual action is required to be inhibited without any ongoing action component.

Here we report results for the extended SIS model (see ^15^, Supplementary Materials for a formal specification of the model). Briefly, the model framework was specified in terms of the possible stimulus/response combinations. Responses could be either no response (NR), left response (LR), right response (RR) or a dual (bimanual) response (DR), whereas the stimulus could be a dual stimulus (i.e., bimanual, DS), a left stimulus (i.e., right stop signal; LS), or a right stimulus (i.e., left stop signal; RS). The structure of the model allows us to not only fit common responses to the different stimuli but also to account for rare events, such as go-omissions (i.e., NR to a DS) and motor errors (e.g., LR or RR to a DS).

We fitted the models in a Bayesian hierarchical manner ^51^ using the default settings in the Dynamic Models of Choice software ^52^, which implements differential evolution Markov chain Monte Carlo to obtain samples from the posterior distribution ^53^. We started by fitting each participant’s data separately. These individual estimates were used as the start points for the hierarchical fitting. Then we ran the hierarchical fitting in an automatic manner until the chains had converged to the posterior using the DMC function **h.run.converge.dmc**. Finally, we ran the chains for 250 more iterations using the DMC function **h.run.dmc**. These final iterations were used for all analyses. Note that before fitting the model, we removed trials in which the initial bimanual responses were not simultaneous (>50ms between left and right button presses, 4% of trials overall) and averaged the left and right button press times as a single response time (RT) measure for bimanual response trials.

To investigate the psychological mechanisms underlying cueing effects, we fit five different versions of the model in which different parameters were allowed to vary according to the cueing condition (i.e., proactive/reactive): **Model1** permitted the dual-go runner to have a different speed in the proactive/reactive conditions; this is consistent with the usual proactive slowing observed in stop signal tasks when participants are made aware of certain aspects of the task, i.e., stopping likelihood or which hand might have to stop in response-selective stopping. Moreover, the speed of both the left- and right-unimanual runners were permitted to vary according to proactive/reactive conditions. Importantly, this model constrained the speed of the dual-stop runner to be invariant across proactive/reactive conditions. The other four models were variants of **Model1**: **Model3** and **Model4** differed from **Model1** in that the dual-go (**Model3**) or selective (**Model4**) runners were not allowed to vary according to the cue conditions. Finally, **Model5** permitted the dual-stop runner to vary according to the proactive/reactive conditions, additionally to the parameters that were allowed to vary in **Model1**. Note that **Model2** is not reported (the nomenclature of model naming comes from the code on the OSF) as this model only considered minor parameter changes based on motor errors (i.e., left response to a right stimulus). We selected the best model according to Ando’s (2010) BPIC criterion which balances model fit and complexity. We provide R code to reproduce all SIS model analyses through the DMC on the Open Science Framework (OSF): www.osf.io/5a2cm.

### Statistical analyses

Generalised linear mixed models (GLMMs) were used to assess behavioural and EMG outcome measures using a Gamma distribution (appropriate for non-negative, positively skewed variables) and identity function, and modelling participants as random factor. Specifically, a GLMM with factors of Condition (proactive, reactive), Trial Type (go, stop), and Success (success, fail) was used for RT and for the onset, amplitude and length of the RT-generating EMG burst. In these models, we allowed for random intercept and random slopes for Condition, Trial Type, and Condition * Trial Type interaction. A similar GLMM, but using 2 factors (Condition, Success), was used for measures derived from stop trials, including SSD, relative RT (i.e., relative to SSD), and relative EMG onset. Measures associated with partial EMG bursts, which were only assessed during successful stop trials, were tested using a GLMM with a single factor of Condition. These measures included the presence of partial burst and the latency of EMG CancelTime. A logistic distribution function was used for the presence of burst, as required for binary variables. All GLMMs were implemented using a full factorial model, i.e., including main effects and interactions. When a significant effect or interaction was found, post-hoc pair-wise contrasts were conducted using Bonferroni correction. Finally, the amount of stopping delay was compared between conditions using a paired t-test.

Additionally, the full time-series of EMG profiles were compared between proactive and reactive conditions using one-dimensional statistical parametric mapping (SPM1d ^33^). Separate tests were performed for each trial type (successful go, failed stop, successful stop with or without a partial EMG burst), and for each synchronisation reference (go signal, SSD, EMG onset of RT-generating burst, and peak EMG).

### Methodological considerations

Our multi-modal approach allowed us to better understand some of the strengths and limitations associated with EMG CancelTime, an important metric of the latency of motor inhibition that has garnered increasing use and support in recent years ^31,35,54^. Foremost, EMG CancelTime provides single-trial measures of stopping latency, whereas non-parametric SSRT estimates based on the race model ^1^ result in a single point-estimate from the entire distributions of RT and SSD.

Moreover, the simple race model assumes there is only one go runner and one stop runner, and hence cannot adequately accommodate response-selective variations where a third (unimanual-go) runner is introduced ^15,55^; or be applied to other experimental tasks, whereas EMG CancelTime can. Second, EMG CancelTime can provide a *direct* measure of the reduction in muscle activity signalling action stopping, and thus more closely reflects the latency of neural inhibitory motor processes than behavioural estimates. For instance, typical latencies of EMG CancelTime coincide well with the time of reduced corticomotor excitability in successful stop trials, around 150 ms after SSD ^8,10^. In contrast, non-parametric behavioural measures of stopping latency are usually estimated indirectly from RT distributions (integration method) or even from point estimates of average go-RT (mean method), both of which inherently over-estimate the stopping latency by virtue of the electromechanical delay, and are also affected by behavioural strategies, such as proactive slowing down of go-RTs ^56^ and response errors, such as trigger failure of the go or stop runners ^55^.

Furthermore, EEG studies found that trials with a partial EMG response showed reduced lateralized readiness potentials and increased frontal negativity ^46,57^. Hence, stratifying stop trials with partial bursts may reveal distinct “neural signatures” (e.g., EEG, MEP, fMRI) compared to trials with no partial response, thus providing an important window to assess the underlying brain mechanisms involved in motor inhibition.

Nevertheless, we also identified sources of bias when using with EMG CancelTime as a measure of stopping latency, which suggests caution should be applied when using and interpretating it. In the Methods section, we highlighted the importance of imposing time and amplitude constraints when identifying partial EMG responses to avoid inclusion of muscle activity unrelated to the task, such as mirror activity or spurious bursts occurring after the behavioural response (stop-RT). Moreover, EMG CancelTime can only be calculated from successful stop trials with a detectable partial response (typically ∼15-50% of successful stops), which may hinder its generalisation. Our simulations suggest it corresponds to a narrow subset of trials where the stop runner beats the go runner by a margin of approximately 90 – 115 ms (details on supplementary material; see also distributions of EMG onsets in Figure 4). In our dataset, this censoring effect caused a down-bias in stop time estimates, which appears to be stronger in the presence of proactive cues. Note that, in theory, the censoring effect on both the lower and upper ends of the distribution could result in either up-or down-bias in stop estimates, depending on which effect is stronger. This censoring may be reduced by adopting strategies that maximise the proportion of stop trials with a partial burst, including (i) the use of stiff buttons ^15,31^, which require more force (and time) to register a response; or (ii) task designs that minimise delays in go-RT, which would favour fast responses during stop trials and avoid a strategy of waiting for the stop signal before responding, such as modifying SSD staircasing to increase stopping success rates and reduce strategic slowing ^58^. In conclusion, EMG CancelTime can be a valuable complementary measure to understand the underlying mechanisms involved in motor inhibition despite its limitations.

## Supporting information

Supplement, Model-based simulations on the effects of trial censoring associated with the use of EMG CancelTime

## Funding

This work was supported by an Australian Research Council Discovery Project grant (DP200101696) awarded to MRH (chief investigator) and DM (overseas partner investigator). AH and DM are supported by a Vidi grant (VI.Vidi.191.091) from the Dutch Research Council (I).

## Author information

### Contributions

S.E.S., Q.F.G., M.R.H, and A.H. wrote the main manuscript text, S.E.S., Q.F.G., and A.H. performed the data analyses, S.E.S. and Q.F.G. prepared the figures. All authors reviewed the manuscript.

## Data availability

All data and replication code are available via the Open Science Framework: www.osf.io/5a2cm.

## Ethics declarations

### Competing interests

The authors declare no competing interests.

## Notes

https://osf.io/5a2cm

### Competing Interest Statement

The authors have declared no competing interest.

