## Supplement, Model-based simulations on the effects of trial censoring associated with the use of EMG CancelTime for "Faster action reprogramming, but not stopping, with proactive cues: Combining EMG and computational modelling in response-selective stop signal tasks"

### Supplementary material: Model-based simulations on the effects of trial censoring associated with the use of EMG CancelTime as a measure of stopping latency

In our main paper, we used the SIS (*simultaneously inhibit and start*) model to fit data from a response-selective version of the stop signal task, performed with or without the presence of visual cues (proactive and reactive conditions). The SIS model describes this task as a race between three processes: Presentation of the go signal triggers (i) the dual-go runner, responsible for causing a bimanual response. Subsequent presentation of the stop signal simultaneously triggers both (ii) the stop runner, in an attempt to inhibit the initial response, and (iii) the unimanual go runner, responsible for causing the new unimanual response. We also presented a detailed methodology used to detect bursts of electromyography (EMG) signals and to estimate EMG CancelTime, which is an EMG-based measure of stopping latency, defined as the latency of the peak of a partial EMG burst relative to the time of the stop signal delay (SSD). Note that, although EMG CancelTime can be directly observable from single-trial EMG profiles, by definition it can only be estimated from successful stop trials where a partial burst can be observed, i.e., it ignores (censors) all trials without an observable partial burst.

Here, we use the parameters from the model that best fit our dataset (**Model 1**) to run simulations that explore the effects of trial censoring when using EMG CancelTime. Using the framework of the SIS model, the presence or absence of a partial EMG response depends on the relative finish times of the stop and dual-go runners (see **Figure S1**):

- Stop runner wins by large margin: Successful stop, no partial burst
- Stop runner wins by small margin: Successful stop with partial burst
- Stop runner barely wins (very small margin): Partial burst cannot be distinguished from the RT-generating burst, i.e., both bursts merge together into a single burst
- Stop runner loses the race: Failed stop

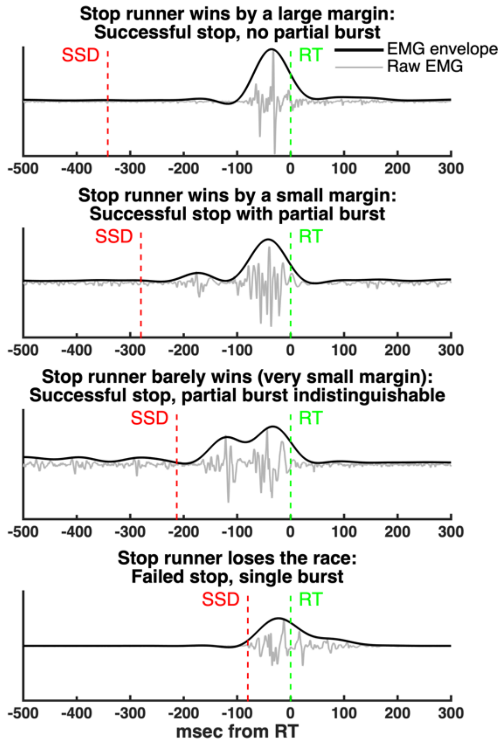


**Figure S1**. Representative examples of single-trial raw EMG signals and the corresponding EMG profiles. The presence or absence of partial EMG bursts depends on the finish times of the dual-go and stop runners. Specifically, a partial burst can only be detected in successful stop trials where the stop runner wins the race by a small margin (second row from the top). As a result, trials where the stop runner is “too fast” or “too slow” are censored, as a partial burst cannot be detected.

Mathematically, if we consider CancelTime to be an EMG-based measure of stopping latency, the corresponding behavioural estimate (i.e., the finishing time of the stop runner) would occur after the electro-mechanical delay (EMD):

$$Stop\ = SSD + CancelTime + EMD$$

Similarly, the finish time of the dual-go runner (i.e., RT) can be expressed in terms of EMG events as approximately:

$$DualGo\ = SSD + CancelTime + \frac{\binom{length}{partial}}{2} + \binom{silent}{period} + \binom{EMG}{incrase} + EMD$$

Hence, the difference between the finish times of the stop and go runners is:

$$[DualGo - Stop] = \frac{(length\ partial)}{2} + (silent\ period) + (EMG\ increase)$$

where:

- Length partial/2: Average time from the peak to the offset of the partial burst (~100/2 = 50 ms)
- Silent period: Time from the offset of the partial burst to the onset of the RT-generating burst (min 20 ms using our EMG detection algorithm)
- EMG increase: Time required to recruit enough motor units to generate the force required to press the button (estimated as at least 20 ms)

To determine the extent of the censoring effect, we estimated the minimum and maximal differences between [DualGo – Stop] where a partial burst can still be detected. The minimum difference (i.e., an "upper threshold" for the finish time of the stop runner) was obtained from the estimations described above as being at least 50 + 20 + 20 = 90 ms. Using this "upper threshold", we simulated different values of the "lower threshold" (maximum difference [DualGo – Stop]) and found that a value of 115 ms resulted in a number of trials included in the simulation that is similar to the proportion of trials with partial burst from our EMG analysis, i.e., ~18% of successful stops, or ~9% of all stop trials.

Using the SIS model, we applied these lower and upper thresholds (90 and 115 ms) to estimate the effects of trial censoring on model-based estimations of stop latency. We found that estimates were down-biased in both the proactive and reactive conditions, although this bias effect was ~7ms stronger in the proactive condition (credible interval[1] of difference: 5 – 9 ms), causing the stop runner to be faster in the proactive condition. These results are consistent with the 8 ms difference in EMG CancelTime between conditions found using GLMMs. To further investigate this censoring effect, we repeated the simulations using a wide range of upper and lower thresholds (see **Figure S2**), and generally found that a down-bias occurred in most cases, although it was sometimes of a smaller magnitude. As would be expected, the effect of censoring bias decreases as the proportion of trials included in the estimate increases (i.e., a larger sample of trials better represents the total). This suggests that strategies that maximise the proportion of trials with partial burst are likely to reduce the censoring effects, although not necessarily in a linear way. Further investigation is required to determine how this effect varies between experimental contexts (e.g., stiff vs. compliant buttons) or behavioural strategies (e.g. strategies that minimise slowing of go-RTs).

[1] Credible intervals were obtained by repeating the calculation for a randomly selected posterior parameter estimates.

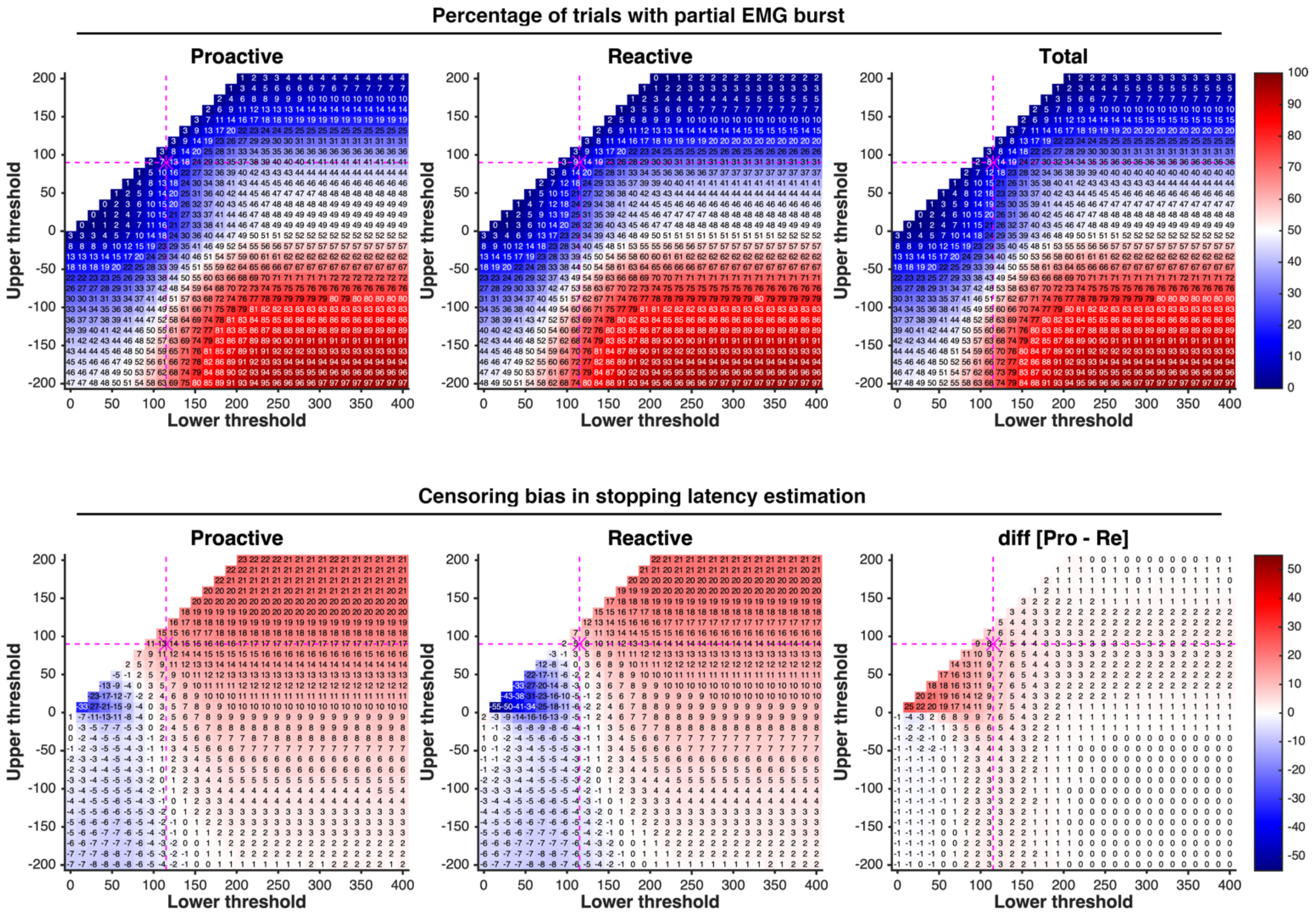


**Figure S2**. Simulated censoring effects on model-based estimates of stopping latency when including only stop trials where the stop runner wins the race against the dual-go runner by a margin that is within the interval between the upper and lower thresholds. Top row: Percentage trials included in the simulation (i.e., percentage trials expected to have a partial EMG burst). Bottom row: Corresponding down-bias in model-based estimates of stopping latency. In all panels, the horizontal and vertical dashed lines (in magenta colour) indicate the thresholds most consistent with our data, and their intersection represents the associated value for the quantity graphed in the panel.
